# PEAK2VEC ENABLES INFERRENCE OF TRANSCRIPTIONAL REGULATION FROM ATAC-SEQ

**DOI:** 10.1101/2021.09.29.462455

**Authors:** Lifan Liang, Xinghua Lu, Songjian Lu

## Abstract

Transcription factor (TF) binding sites in ATAC-seq are typically determined by footprint analysis. However, the performance of footprint analysis remains unsatisfying and most TFs do not exhibit footprint patterns. In this study, we modified the convolutional neural network to project sequences into an embedding space. Sequences with similar nucleotide patterns will stay close together in the embedding. The dimensionality of this embedding space represents binding specificities of various TFs. In the simulation experiment, peak2vec accurately distinguished the three TFs in the embedding space while conventional deep learning cannot. When applied to the ATAC-seq profiles of hepatitis carcinoma, peak2vec recovered multiple motifs curated in database, while significant portion of sequences corresponding to the TF are located at the promoter region of its regulated genes.

## INTRODUCTION

Since first described in 2013 (1), ATAC-seq has gained particular popularity. The exponential increase of ATAC-seq curated datasets indicates its value in a wide spectrum of biological studies (2). In particular, the project of TCGA has profiled 410 tumor samples with ATAC-seq to interrogate the transcriptional regulation (3). Unlike TF ChIP-seq, the accessible genome contains all the sequences potentially interacting with transcription factors, offering a great opportunity to systematically analyse gene regulation.

Currently, the major approach to analyse TF activities in ATAC-seq is to utilize the footprint pattern formed by the cleavage signals (4, 5). Although these methods have yielded insights into transcriptional regulation, only one fifth of TF motifs show protection from the DNA cleavage (6). Relying on footprinting analysis may severely hinder the investigation into transcriptional regulatory mechanism. Other types of information, especially DNA sequences highly enriched in the assay, may serve as important alterative clues to understand transcriptional regulation.

Computational researches have utilized the sequence information to complement the footprint pattern, such as TFEA (7) and MEME-centrimo (8). TFEA performed TF enrichment analysis on the peak regions in the differential analysis to identify TFs causally responsible for biological differences between samples. MEME-centrimo compared the cleavage counts of predicted binding sites with the cleavage events around them. However, these methods relied on curated TF motifs, which may be incomplete, inaccurate, and inconsistent across tissues and conditions.

On the other hand, deep learning has been successfully applied to de novo identification of TF motifs in TF ChIP-seq experiments, with the best performance among different approaches (9).

However, despite various improvements (10–12) and expanded application (13, 14) along the past five years (15), we have rarely seen any deep learning applications to identify TF binding activities in the chromatin accessibility profiles. This is probably because conventional deep learning algorithm may not be a suitable tool when there are multiple TFs mixed in the set of sequences. This will be demonstrated in the simulation experiment.

In this study, we presented a novel variant of convolutional neural network (CNN), Peak2vec, to perform de novo inference of the coregulation among enriched regions by constructing the embedding space from the peak sequences in ATAC-seq. We hypothesized that by modifying existing deep learning algorithm, it is possible to uncover various TF binding specificities and downstream regulon within the chromatin accessibility profiles without relying on current knowledge.

## MATERIAL AND METHODS

The overall architecture of Peak2Vec is illustrated in Figure 1. For each fix-length DNA sequences, we generated the one-hot encoding matrix for both strands. The binary matrix is scanned by multinomial convolution of different sizes. Normalized convolution output is concatenated together as the embedding vector. The kernel parameters and the embedding space of peak sequences is learned by classifying whether a sequence was a peak region or a random sequence. Features with positive correlation to the real peak regions were selected as the embedding vector for enriched sequences.

**Figure 1.**
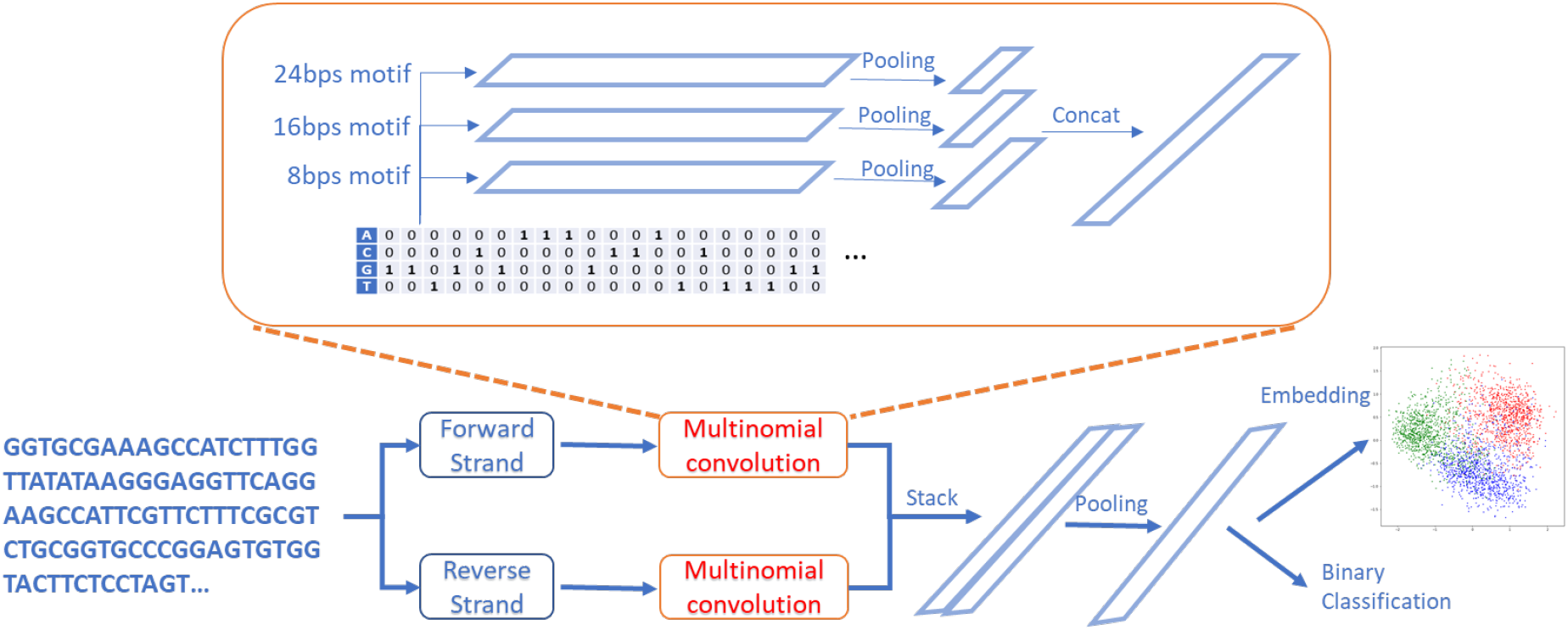
The architecture and workflow of Peak2vec. DNA sequences are transformed into one-hot encoding array. Three layers of multinomial convolution are applied to the array. Three max pooling layer are applied to each convolution output to generate feature representation that are concatenated for binary classification. After training the model for binary classification. Features with positive coefficients towards peak region prediction were selected to construct the embedding space of peak sequences.

### Multinomial convolution kernel

In this section, we show how 1D convolution connects to multivariate multinomial mixture in theory and introduce the basic convolution unit in Peak2Vec. First of all, 1D convolution is basically applying a sliding window (named “kernel”) over an “image” along the first dimension. Suppose the input “image”, *C*, is an N by M matrix, the kernel, *W*, is an N by K matrix, then the output (named “feature map”), *C*\*, would be a vector of length (*M* – *K* + 1). The relationship between the input and output is:

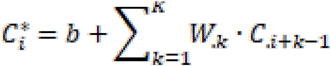

where *W_.k_* · *C*_.*k*+*i*−*1*_ is the inner product between the *k*th column of W and the *k*+*i*−1th column of *C, b* is scaler for bias terms. This is the same one-dimensional convolution as in DeepBind or DeepSEA. Usually, multiple sliding windows will be applied to the “image”, hence the feature map, *C*\*, expands to a matrix of L by (*M* – *K* + 1), given that L is the number of kernels. And the bias term, *b*, expands to a vector of length L.

However, in this study, we apply the Softmax function to each column of W such that each column sums up to one. In this way, the *l*th kernel, *W_l_*, can be interpreted as the parameterization of a multivariate multinomial distribution *Z_l_*. Hence *l*th component of multinomial distribution Z, the *k*th column of W, *W_.k_*, represents that

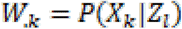

In addition, we also perform Softmax on the bias term. So *b_l_* represent the prior probability of *Z_l_*. Furthermore, we take the log of *W* and *b* elementwise before applying convolution. Considering the computation of 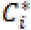, we have

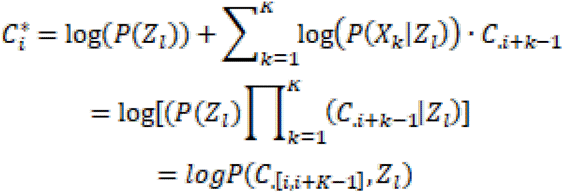

The output C* would become the log joint probability of the i to i+K-1 columns of C and the *l*th kernel. Then we further perform the Softmax function over each column of C*. Clearly, 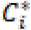 after Softmax dictates the posterior probability of *Z_l_*.

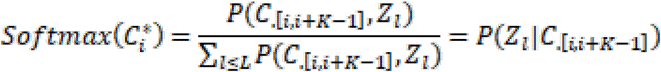

At this point, it is clear that the specialized convolution kernel here is analogous to perform the EM algorithm for multivariate multinomial mixture on each sliding window. The forward computation is inferring the membership of samples, the E step. The backward propagation is learning distribution parameters and priors, the M step. This is only an analogy because of two major differences: (1) we do not identify the maximum likelihood in each iteration of backpropagation as in the real M step; (2) the distribution parameters are shared across all sliding windows. Still, this theoretical connection may provoke insights in theoretical analysis and deep learning models development.

In summary, we applied log Softmax function to the kernel weights and biases before convolution. After convolution, we applied Softmax to the feature map. In this way, kernel weights (W) and biases (b) can be interpreted as components of multivariate multinomial mixture and corresponding priors. The feature map can be regarded as the posterior probability that the scanned section of C is sampled from the *l*th kernel.

In addition, as shown in Fig 1, feature map of the first and the second convolution layer directly feed to the next convolution layer without pooling. This can reserve position information for the next convolution. However, only the first convolution layer is interpretable because it is directly connected to the one-hot encoding matrix of DNA sequence. Although the other convolution layers cannot be interpreted as position weight matrix (PWM), they enable the model to capture complicated regulatory sequence patterns.

### Max pooling layer

The max pooling layer outputs the maximum value of every row of *C*\*. This means that a kernel is activated if it has high posterior probability in any subsequence of the sample. Output of this layer become features for the classification task. The feature size is the number of kernels in the corresponding convolution layer.

### Model training

All the outputs from pooling layers are concatenated as an embedding vector U. The embedding was train by binary classification on whether the sequence contains regulatory elements or not. Suppose the binary label is a vector Y of length S (sample size), then the objective is the cross entropy between Y and the estimated Y.

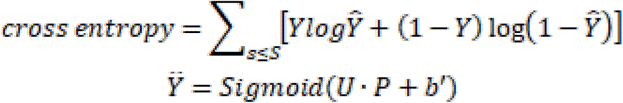

where Sigmoid is the sigmoid function, P is the weights, b’ is the bias term. The model is trained with gradient-based optimization, specifically RMSprop.

### Handling the reverse complement strand

The reverse complement also went through the variable length multinomial convolution and produce a concatenated embedding vector. As shown in Figure 1, we selected the bigger value from both strands for each feature and constructed the final embedding vector for training.

### Data preprocessing

The set of positive sequences were constructed by extending certain length upstream and downstream to the peak summits of ATAC-seq. In simulation experiment, the extension length was 100 base pairs. In real data analysis, the extension length was 250. We generated negative sequence with the same dinucleotide frequency from the positive sequence. A sequence of length M is then transformed into 4 by M matrix via one-hot encoding.

### Embedding vector interpretation

We used Gaussian mixture to identify the clusters of the embedding vectors. For each cluster, samples are divided into two groups, samples in the cluster and samples in other clusters. We then perform Wilcoxon test to identify motif features with significantly higher values in the cluster than outside the cluster. The feature with the smallest p value is regarded as the signature motif for the corresponding cluster.

In addition to the cluster analysis, we cross examined the convolution kernels with motif matching by TomTom (16) and gene set enrichment analysis by ChEA3 (17). Kernels matched to the same TF were selected to complement the clustering analysis. For the gene set enrichment analysis, we selected 1000 sequences with the top values in each kernel and annotated to genes. The set of gene symbols were the input to ChEA3.

## RESULTS

### Peak2vec recovers gene regulation accurately

To validate Peak2Vec’s capability of identifying different structural regularities from enriched DNA sequences, we collected TF ChIP-seq of JUN, CTCF, and POLR2A from the HepG2 cell line in the ENCODE project. We selected the top 1000 peaks from each TF in terms of q values. 6000 negative sequences were also generated. Peak2Vec was trained on 9000 samples in total. Peak2vec has three layers of multinomial convolution. Each layer has 256 kernels.

We then perform conventional clustering algorithms on the embedding space. As shown in Figure 2, the three TF can be identified in different clusters despite the clustering algorithm. In subsequent real data analysis, we continued to use Gaussian mixture to identify sequences attached to different TFs in the simulation dataset.

**Figure 2.**
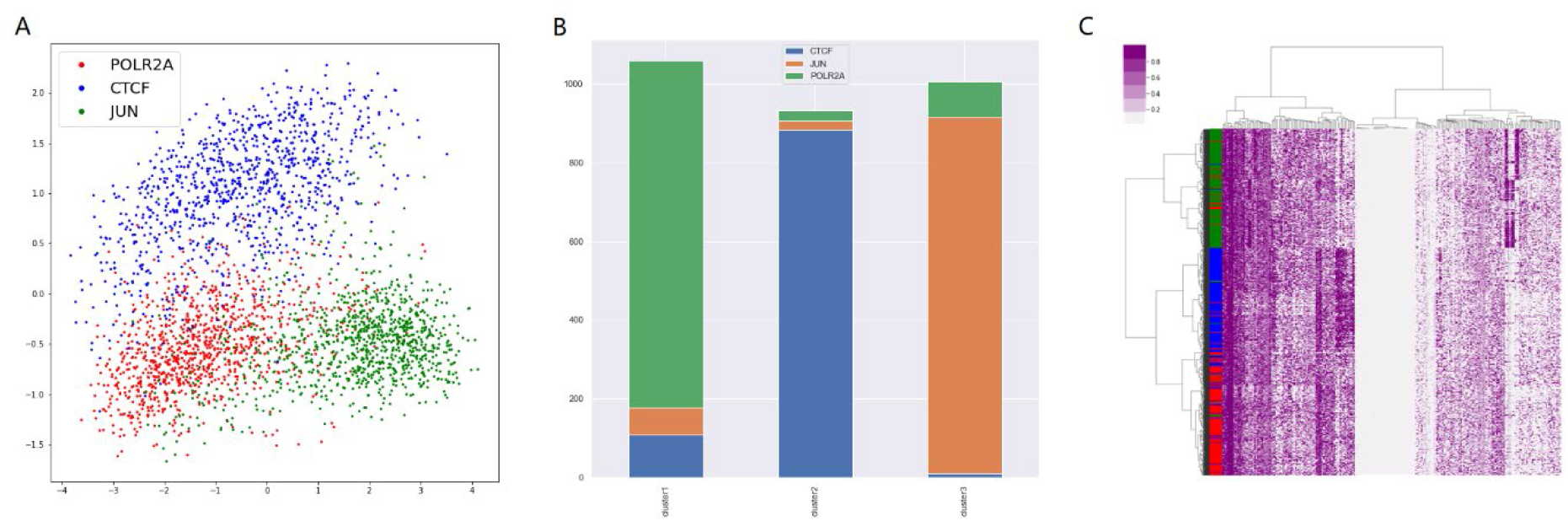
Different methods exploring the embedding space of peak2vec. PCA visualization of the embedding space (A), Gaussian mixture (B), and hierarchical clustering (C) of the embedding vectors.

To demonstrate the necessity of multinomial convolution, we also implemented a convolutional neural network (CNN) model with conventional convolutional kernels and ReLu activation as described in DeepBind (9). Other than the kernels, our implemented CNN has the same architecture and the same training procedure as the peak2vec. The results are shown in Figure 3. Clearly, CNN fails to distinguish different TFs in the embedding space.

**Figure 3.**
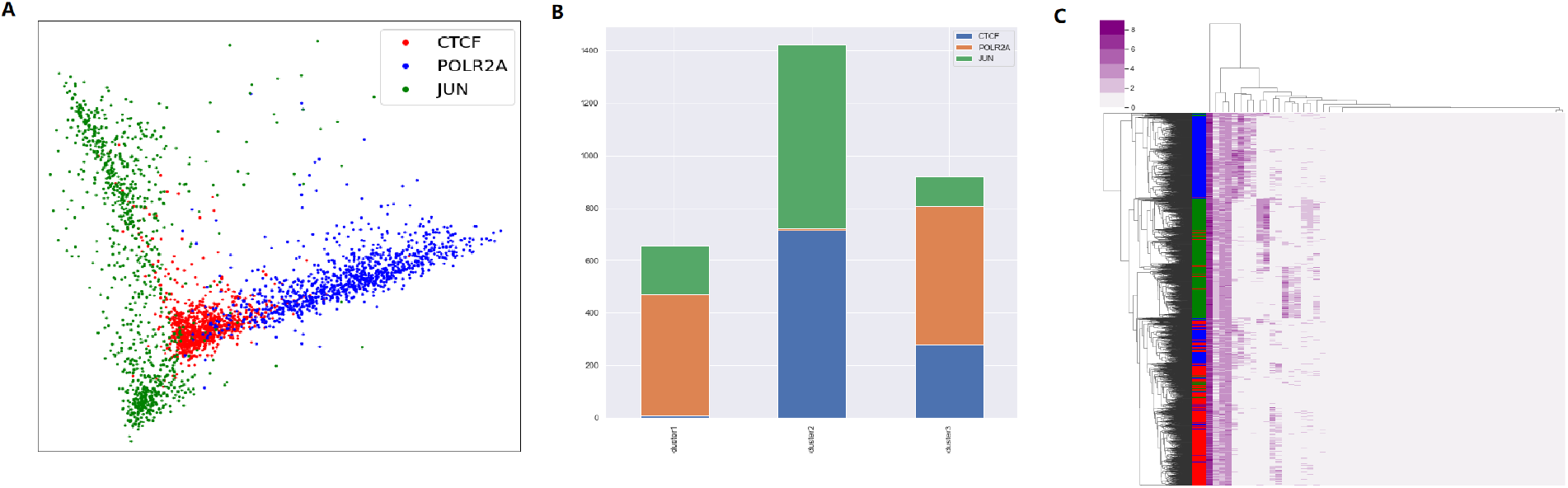
Different methods exploring the embedding space generated by conventional convolutional neural network (CNN). PCA visualization of the embedding space (A), Gaussian mixture (B), and hierarchical clustering (C) of the embedding vectors.

We further extract the signature motif for the three cluster. As shown in Figure 4, the motif signature is similar to the motif in JASPAR. In the case of CTCF, “CCTCC” is similar to “CCACC” in JASPAR. As for JUN, the first four base pairs match nicely to the motifs in JASPAR. We cannot find motifs for POLR2A in any database. JASPAR also cannot find any high-quality motifs for POLR2A.

**Figure 4.**
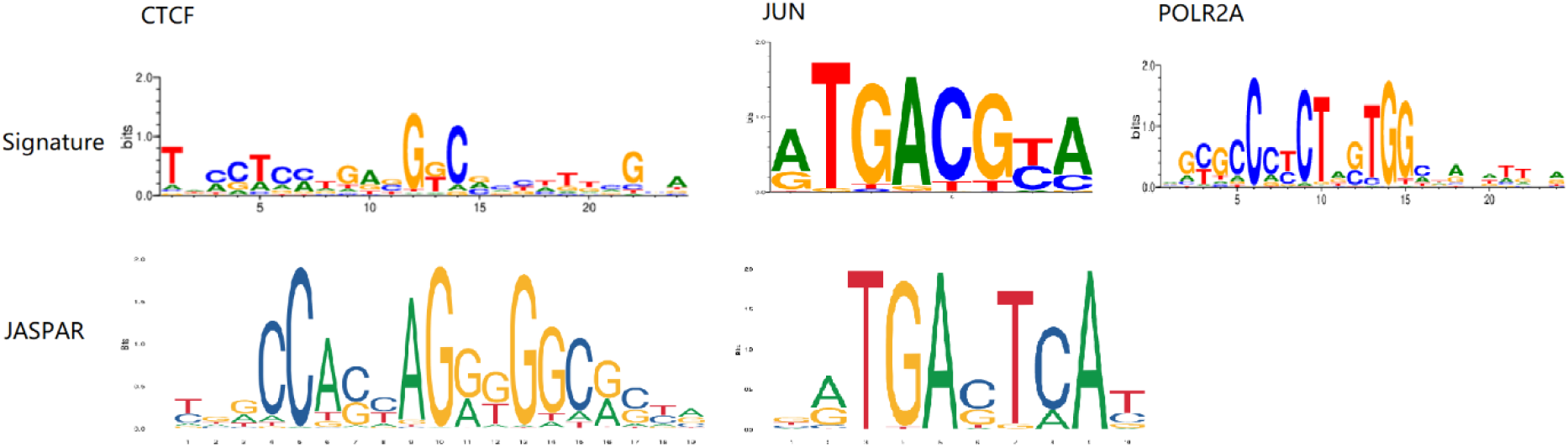
Comparison between signature motif from Peak2vec (above) and motifs from JASPAR (below). There are no curated motifs for POLR2A.

Still, Figure 2 showed that the embedding of POLR2A peak sequences are distinguishable from the other two.

### Application to ATAC-seq profiles reveals regulation of multiple TFs

We downloaded liver cancer type-specific peak signals from Xena Browser. 26513 peaks with signal values above 10 have been selected. All the sequences have a fixed length of 500 bps. We then generated 53026 negative sequences following the same dinucleotide frequency of the peaks. The model is significantly larger than that in simulation experiment, with 1024 kernels in each layer.

After extracting the embedding vectors from Peak2vec, we conducted Gaussian mixture on the embedding vectors. The number of clusters was set to 80, as determined by the Akaike information criterion (AIC). The signature motifs were extracted to compare with the motifs curated in JASPAR vertebrate dataset. Signatures extracted from clusters seems to be redundant for many unique clusters. As shown in Figure 5, the signature motif of some clusters can be matched to known motifs quite nicely.

**Figure 5.**
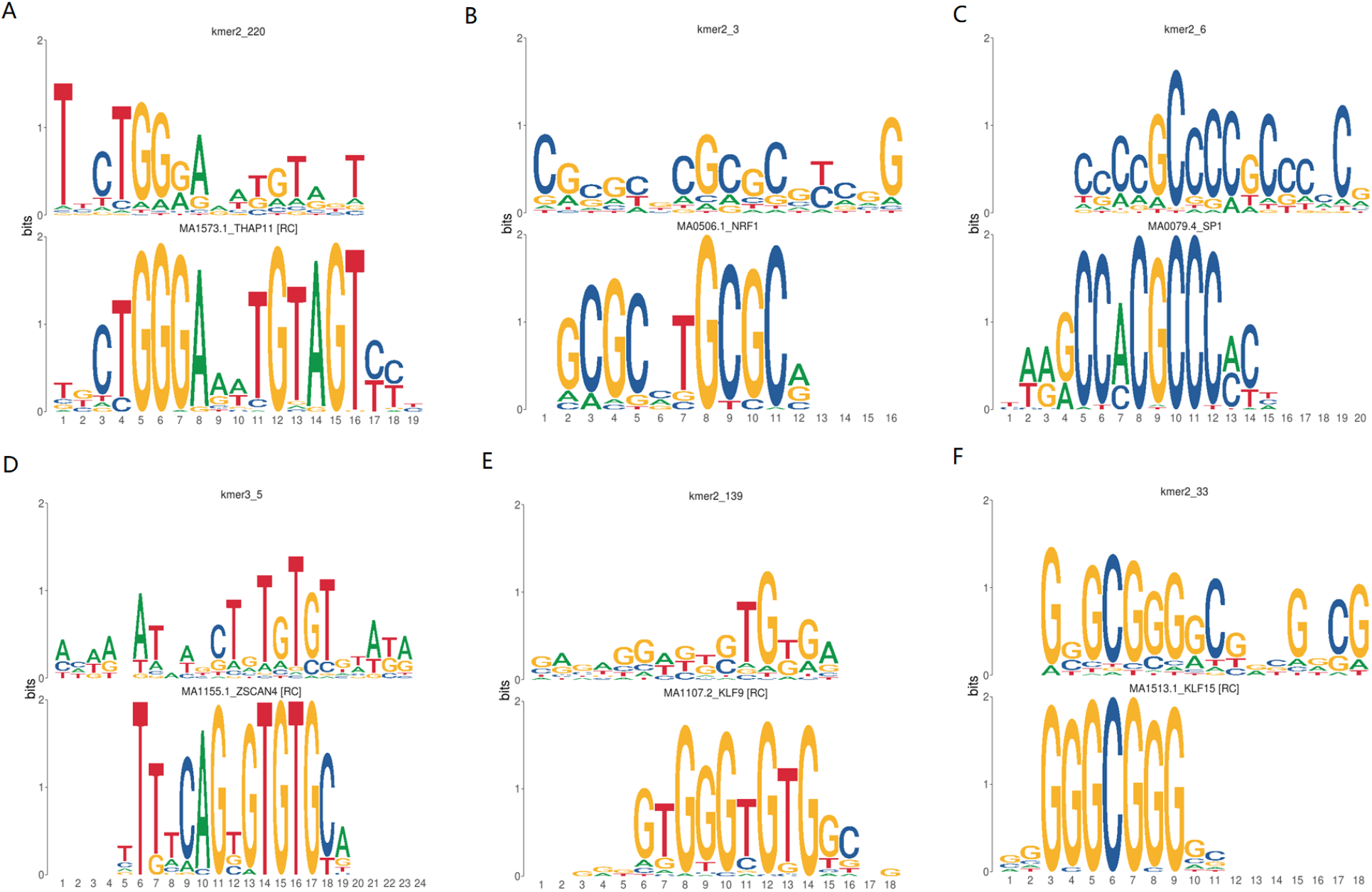
Signature motifs that can be matched to known motif in JASPAR. For each subplot, the upper part is the signature and the bottom part is the known motif. (A) is matched to the transcription THAP11; (B) is matched to the transcription factor NRF1; (C) is matched to SP1; (D) is matched to ZSCAN4; (E) is matched to KLF9; (F) is matched to KLF15.

In addition, we search through the multinomial convolution kernels other than cluster signatures to identify meaningful motifs. The search is taking the intersection of motif matching results and gene set enrichment analysis results. Motif matching is performed in the same way as above. As for gene set enrichment analysis, we extracted peaks with top 1000 score for each kernel and feed them to ChEA3 for enrichment analysis. As shown in Figure 6, brute force searching also reveals other motifs missed by the clustering analysis. These motifs have been matched significantly to both JASPAR and the gene set enrichment analysis.

**Figure 6.**
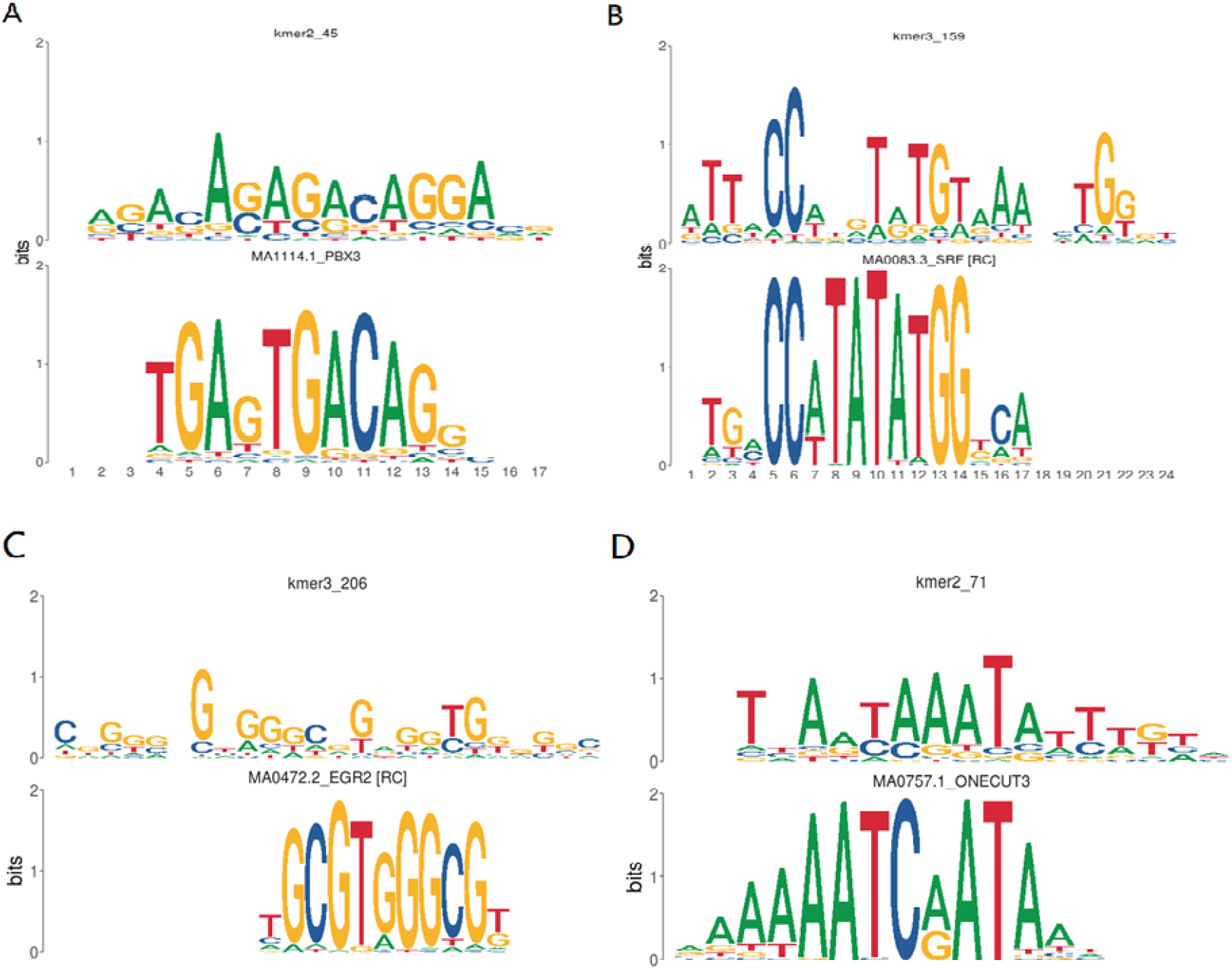
Motif identified through searching all the kernels. (A) was matched to PBX3 with enrichment analysis FDR=5.37e-6; (B) was matched to SRF with ENCODE enrichment analysis FDR=0.014; (C) was matched to EGR2 with ENCODE enrichment analysis FDR=0.0475; (D) was matched to ONECUT family with ReMAP enrichment analysis FDR=8.34e-6.

We also found that although some convolution kernels have not found a significant match to the curated motifs, they may still be biologically meaningful. Examples in Figure 7 showed that some kernels may capture a pair of TFs at work (Figure 7A) or not distinguishable enough (Figure 7B).

**Figure 7.**
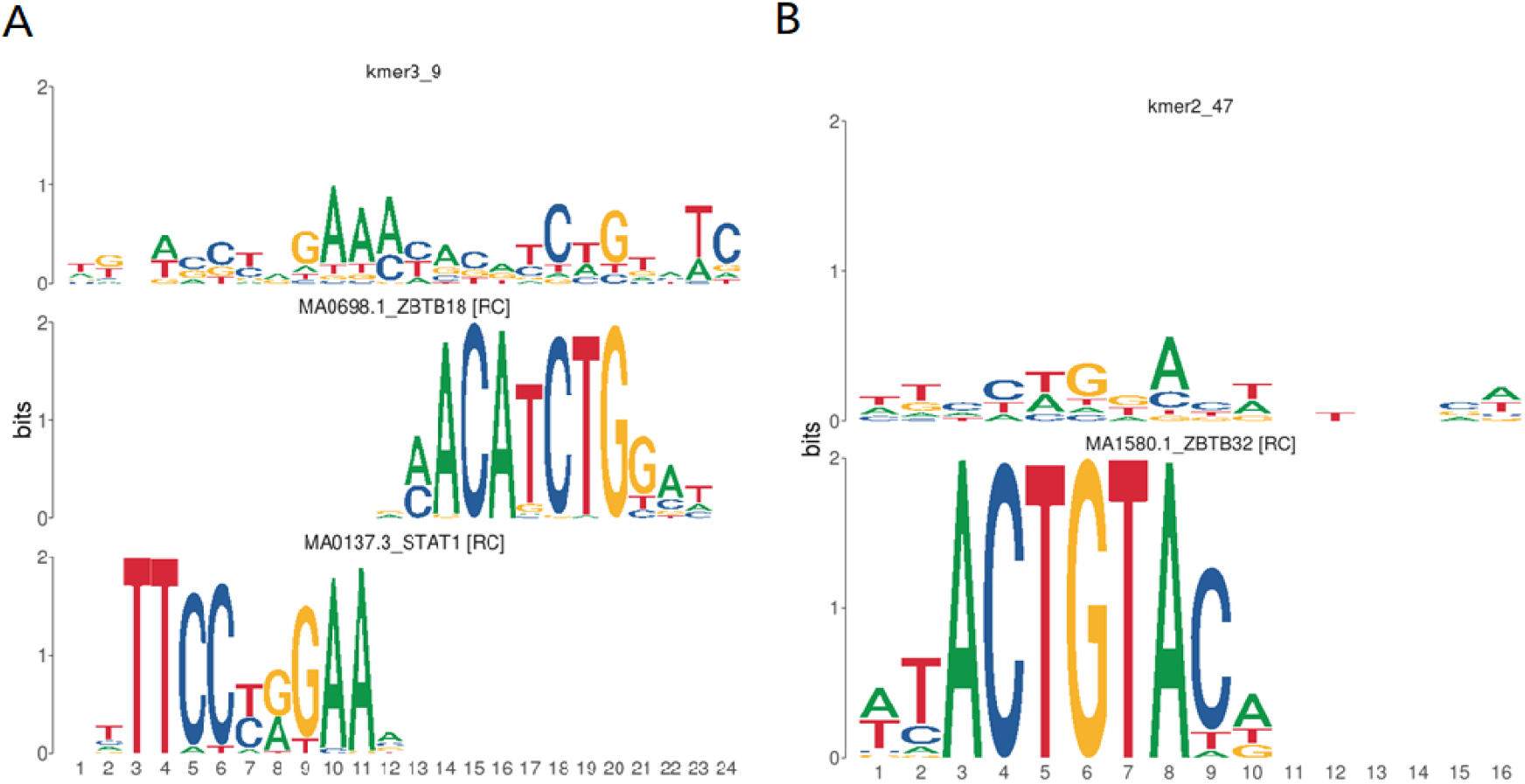
Learned motifs that only found a weak match in the JASPAR database. Kernel in (A) consists of two known TF motifs, ZBTB18 and STAT1. Kernel in (B) can be matched to ZBTB32 if only the top nucleotide was considered.

## DISCUSSION

In this study we presented Peak2vec, a novel algorithm that can identify ATAC-seq peaks regulated with the same TF, while providing the corresponding signature motif. To our knowledge, this is the first model to infer transcriptional regulation relying solely on ATAC-seq. The idea of employing similar deep learning algorithms to study epigenomics has been quite popular along the past decade (9, 14, 18). We contributed to the methodology by adapting the convolution kernels so that it can identify multiple TFs within the chromatin accessibility profiles. Our results show that peak2vec is effective when handing with sequences with diverse binding specificity.

Peak2vec is also easier to interpret since a multinomial convolution kernel directly represents a position weight matrix (PWM). When applying deep learning to extract motifs from TF ChIP-seq, previous research (9) need to align the sequences according to activated position in the feature map.

Still, there is much room for improvement. One particular issue is that enrichment analysis for coexpression module did not overlap with motif comparison much. It is probably due to two reasons: (1) peaks were mapped to the nearest genes. This may not be accurate especially for peaks in the remote regions. They may be trans-regulatory elements; (2) TFs may have different binding sites in different tissues. In the future, we need to utilize the RNA expression information to determine the correlation between peaks and genes, which has been partially employed in the TCGA project (3).

Another limitation is that although Peak2vec is capable of finding novel TF motif, it is difficult to determine the identity of the TF whose motif are not curated. Future research may develop methods to ascertain TF identity given the motif and corresponding regulon inferred from Peak2vec.

In addition, we performed Gaussian mixture on the embedding vector. This method performed well when the number of clusters is known. Such is the case with the simulation experiment. However, in real data experiment, the number of TFs involved are unknown. When determined by AIC, the number tend to be too large and led redundant clusters. In the future, we need to refine the workflow on how to extract the TF motifs and its corresponding regulon.

Pease note that application of peak2vec is not restricted to ATAC-seq. Any set of fixed size sequences can be the input. For example, peak2vec may also be applied to TF ChIP-seq experiment in case multiple motifs exists for cofactors. In the future, we may expand the application of peak2vec to other types of high-throughput technologies.

This study presented a novel algorithm named peak2vec. We believe peak2vec would serve as a valuable tool for biologist to analyse transcriptional regulation from ATAC-seq profiles. It also deserves attention of computational researchers due to its general utility to extract sequence information from sequencing data.

## DATA AVAILABILITY

Genome Maps is an open source collaborative initiative available in the GitHub repository (https://github.com/compbio-bigdata-viz/genome-maps)

Atomic coordinates and structure factors for the reported crystal structures have been deposited with the Protein Data bank under accession number XXXX.

## ACKNOWLEDGEMENT

The results <published or shown> here are in whole or part based upon data generated by the TCGA Research Network: https://www.cancer.gov/tcga.

